# Reduced body size of *Varroa destructor* associated with varroa-resistant honey bee colonies across Europe

**DOI:** 10.64898/2026.03.11.711027

**Authors:** Adéla Krajdlová, Václav Krištůfek, Alena Krejčí

**Affiliations:** University of South Bohemia, Faculty of Science, 370 05 Ceske Budejovice, Czech Republic; Biology Centre, Institute of Entomology, Czech Academy of Sciences, 370 05 Ceske Budejovice, Czech Republic; Biology Centre, Institute of Soil Biology and Biochemistry, Czech Academy of Sciences, 370 05 Ceske Budejovice, Czech Republic

**Keywords:** *Varroa destructor*, *Apis mellifera*, mite morphology, varroa-resistance

## Abstract

The ectoparasitic mite *Varroa destructor* is the most significant parasite of the Western honey bee (*Apis mellifera*) and a major driver of colony losses worldwide. Although extensive research has focused on behavioral and physiological mechanisms of host resistance, comparatively little attention has been paid to potential phenotypic responses of the parasite itself. Here we investigated body size variation in *Varroa destructor* associated with varroa-resistant and non-resistant honey bee colonies across four European countries. We quantified the dorsal shield area of adult female mites from multiple colonies differing in the honey bee colonies resistance status, using standardized digital image analysis. Across geographically distant non-resistant populations, mite body size was remarkably consistent, with a median dorsal shield area of 1.47 mm^2^. In contrast, mites originating from varroa-resistant colonies were consistently smaller, with a median dorsal shield area of 1.37 mm^2^, representing an approximately 6.8% reduction in body size. This pattern was reproducible across different geographical areas, honey bee genetic backgrounds and beekeeping practices. The striking stability of mite body size in non-resistant populations contrasted with the consistent reduction observed in mites associated with resistant hosts, suggesting a host associated shift in parasite phenotype. Because body size in arthropods integrates developmental conditions, nutritional availability and resource allocation, the observed pattern may reflect altered developmental environments and selective pressures imposed by resistant hosts. Our results show a consistent morphological shift in this globally important parasite associated with resistant hosts and suggest that dorsal shield size in *Varroa* could serve as a new selection marker for varroa-resistant honey bee colonies.

## INTRODUCTION

The ectoparasitic mite *Varroa destructor* is regarded as the most significant biotic threat to the Western honey bee (*Apis mellifera*), contributing to colony losses through direct parasitism and the transmission of viral pathogens, particularly Deformed wing virus (DWV) [1; 2; 3]. Since its host shift from the Asian honey bee (*Apis cerana)* and subsequent global spread, the mite has fundamentally affected *Apis mellifera* colonies survival particularly in the northern hemisphere and remains a primary factor limiting sustainable apiculture [4; 5]. As a consequence, this host parasite system has become a good model for studying the dynamics of parasite virulence, host resistance and disease transmission in managed animal populations.

In response to the negative impact of *Varroa* parasitation on honey bee colonies health and survival, increasing attention has been directed toward honey bee colonies that survive mite infestation without regular chemical miticide treatment [6; 7]. They are associated with host traits that reduce the mite reproductive success, including varroa-sensitive hygiene, grooming behavior and brood cell recapping, supporting better colony survival [8; 9; 10; 11; 12]. Varroa-resistant populations have been described across diverse geographic regions [13] and they were established either through natural selection or through dedicated breeding programmes that evaluate indicators of varroa resistance in individual colonies and propagate queens from the best performing colonies. However, selection based on assessing mite fertility in honey bee brood, hygienic behavior or the rate of brood cell recapping is time consuming and the heritability of these traits is relatively low. Identifying additional selection marker that can be incorporated into the breeding programmes would therefore represent a significant advancement.

While most studies have focused on behavioral resistance mechanisms in the host and the reproductive parameters of the mite [14; 15; 16], comparatively little attention has been given to morphological characteristics of *Varroa destructor* [17; 18; 19]. Body size in arthropods represent integrative outcomes of developmental conditions, nutritional availability and resource allocation and it may influence key parameters such as fecundity, feeding efficiency and survival in arthropods [20; 21]. Because *Varroa destructor* completes its reproductive cycle within capped brood cells of honey bee, developmental environment determined by host traits may influence mite phenotype and reproductive success. Studying morphological variation of the mite across populations could therefore provide insights into how host imposed conditions influence parasite phenotype. Moreover, such morphological changes may be used as new markers for targeted selection of varroa-resistant honey bee colonies. Despite the potential biological relevance and the global importance of honey bees, comparative assessments of morphological variation in *Varroa* associated with host resistant traits remain largely unexplored.

In the present study, we analyzed body size variation in *Varroa destructor* collected from varroa-resistant and non-resistant honey bee colonies in four European countries. We hypothesized that mites associated with resistant colonies differ in body size compared with mites from non-resistant colonies, reflecting the parasites adaptive responses to host imposed selection pressure. Integrating samples from diverse geographic regions allowed us to determine whether observed differences are consistent across different environmental conditions, host genetic origin and colony management regime. Detecting a significant change in mite size in varroa-resistant colonies could provide a practical new marker of varroa resistance.

## METHODS

### Apiaries involved in the study

The GPS locations of all apiaries included in the study, along with the beekeepers names and dates of mite collections are listed in Supplementary table ST1. In all apiaries, several colonies were examined and mites from each colony were analyzed separately, except for samples labeled ‘m’ that consisted of pooled debris from multiple colonies from the same apiary.

#### Czechia

Mite samples were collected from fifteen non-resistant treated apiaries and one varroa-resistant nontreated apiary. All apiaries consisted of *A*.*m*.*carnica* colonies.

The non-resistant colonies were kept in Czech national hives (frames 390×240 mm) or Langstroth 2/3 hives (frames 482×159 mm). They were treated with formic acid or flumethrin in summer, amitraz fumigation in autumn and in some cases also oxalic acid trickling in winter. Wax foundations were used for building combs. Honey was harvested in June and July and supplementary feeding with sucrose syrup was provided in August (typically 15–20 kg per colony). The average winter losses in these treated non-resistant apiaries fall within the range of 10-20%.

The varroa-resistant colonies were kept in Langstroth 2/3 hives (frames 482×159 mm). They were not exposed to any chemical miticide treatment since 2015. Only foundation-free comb building was allowed in these colonies. Varroa management included open-brood trapping comb for newly established splits and total brood removal and queen exchange in colonies with high varroa or virus infestation. Queen breeding in this apiary was based on selection of genetic material from colonies with long term winter survival, low varroa mite numbers in debris and long term absence of adult bees with visible signs of viral infections. Honey was harvested in June or July and supplementary feeding with inverted sugar syrup was provided in July (typically 15–20 kg per colony). The average winter losses in this apiary fall within the range of 10-20%.

#### United Kingdom

Mite samples were collected from three non-resistant treated apiaries and one varroa-resistant nontreated apiary in Wales. The non-resistant apiaries consisted of *A*.*m*.*mellifera* and *Buckfast* colonies, the genetic origin of the varroa-resistant bees is uncertain but *A*.*m. mellifera* and *A*.*m. caucasia* are the most prevalent bee races in Wales [22].

The non-resistant colonies were kept in British national hives. They were treated with thymol (Apiguard) in late summer and oxalic acid in winter. Commercially available wax foundations were used for building combs. Honey was harvested in August or September and 12.5 kg of sugar candy or 10 kg sugar syrup was provided for winter. The average winter losses in these apiaries are 35%.

The varroa-resistant apiary was originally set from local caught natural swarms and no varroa treatment was applied since 2007. The bees were kept in Warre hives and in British national hive. Only foundation-free comb building was allowed in all of the colonies. No queen replacements was done and the life of colonies dated from introducing the swarm to becoming queenless (not during the course of this study). New swarm would then be settled into a refurbished hive. Some colonies were aged over 10 years. Honey was harvested in August/September, leaving enough for winter. No supplementary feed was offered. The average winter losses in this apiary are 14%.

#### Sweden

Mite samples were collected from one non-resistant treated apiary and one varroa-resistant nontreated apiary. The non-resistant apiary consisted of *A*.*m*.*mellifera* colonies, the varroa-resistant apiary kept Elgon bee line (breed of *A*.*m*.*monticola, A*.*m. carnica* and *A*.*m. ligustica* [23]).

The non-resistant colonies were kept in Swedish standard hives (frames 366 x 222 mm) They were treated with oxalic acid trickling during broodless winter period. Wax foundations of regular cell sizes were used for building combs. Honey was harvested in June and July and supplementary feeding with sucrose syrup was provided in August (typically 10 kg per colony). The average winter losses in this apiary are up to 20%.

The varroa-resistant colonies were kept in single walled fir tree boxes with frames 448×137mm. They were not treated against varroa since 2016. They were maintained on small size combs (4.9 mm), using wax foundations. Honey was harvested in June and July. About 15 kg of honey was left for the colony winter times and additional feeding with sucrose syrup was provided in August (typically 10 kg per colony). The average winter losses in this apiary are 5-15%.

#### France

Mite samples were collected from one varroa-resistant nontreated apiary, six varroa-resistant treated apiaries and one treated apiary close to two varroa-resistant apiaries. The varroa-resistant apiaries kept honey bees that were a mix of races resulting from selective breeding based on low varroa reproductive success and colony survival [24]. All analyzed colonies were kept in Langstroth hives (combination of deep brood box with 1/1 frames 482×232mm and shallow supers 2/3 frames 482×159mm).

The varroa-resistant colonies of John Kefuss (group 1) were not treated against varroa since 1999. Foundations from own wax were used for comb building. Honey was harvested in July and 5kg of supplementary sugar candy was provided as winter feed. The average winter losses in this apiary in the season of 2022/23 were 8,11%.

The varroa-resistant colonies of Cyril Kefuss (group 2) used queens from the varroa-resistant population established by John Kefuss but the colonies were treated with amitraz (Apivar) in the autumn. Foundations from own wax were used for comb building. Honey was harvested between May and August and 5 kg of supplementary sugar candy was provided as winter feed. The average winter losses in this apiary in the season of 2022/23 were 9,6%.

The colonies of Victor Kohut (group 3) were located 1.8 km far from one of John Kefuss’s varroa-resistant apiary and 3.72 km far from one of Cyril Kefuss’s varroa-resistant apiary. They were treated with amitraz (Apivar) and used commercially available wax foundations. Honey was harvested between April and August and supplementary sugar candy was provided from September to March when needed. The approximate winter losses in this apiary in the season of 2022/23 were 10-15%.

### Varroa sample collection

The mites were collected from debris on the hive floor following the insertion of a plastic mat on the bottom board. The dates of debris collections were similar between the non-resistant and varroa-resistant apiaries within the same country. Samples were frozen at −20 °C for several hours to eliminate potential damage of the debris by wax moths, then either transferred immediately to the laboratory or sent by post. The collection dates of individual samples are provided in Supplementary table ST1. All mites were carefully separated from the debris using a fine brush and a random subset of adult mature females was selected for slide mounting and subsequent image analysis.

### Measurement of varroa size

A random subset of 23 adult female mites from each sample was mounted on a microscope slide in 50% glycerol and covered with a coverslip, avoiding any bubbles. Images were acquired using Olympus SZX12 microscope at a resolution of 1360 × 1024 pixels. Image analysis was performed in ImageJ software [25], where the surface area of each *Varroa* dorsal shield was measured from a dorsal (top) view. Fewer than 23 mites were mounted in some samples from varroa-resistant colonies due to limited mite availability.

### Statistics

Comparison of mite sizes from varroa-resistant and non-resistant colonies within a country was assessed by Students’s t-test and showed p < 0.001 for Czechia, UK and Sweden.

The effect of colony resistance status on mite size was determined by analysis of complete data sets with all colonies from all countries, using a linear mixed-effects model with colony as a random factor. Mites from resistant colonies were significantly smaller than those from susceptible colonies (likelihood ratio test: χ^2^ = 195.7, df = 1, p < 0.001).

To assess whether this effect was consistent across countries, mite size was further analysed after excluding France dataset due to the uncertain status of control colonies. A linear mixed-effects model with colony as a random factor and country as a fixed effect confirmed that mites from resistant colonies were significantly smaller than those from susceptible colonies (estimate = −0.065 ± 0.008 SE, p < 0.001). The effect of resistance status on mite size did not differ among countries (interaction test: χ^2^ = 2.05, df = 2, p = 0.36).

The effect of varroa treatment on mite size was tested using a linear mixed-effects model including colony as a random factor. Varroa treatment was unevenly distributed among countries (χ^2^ = 39.99, df = 2, p < 0.001), therefore country was included as a fixed effect in the statistical models. Miticide treatment did not significantly affect mite size after accounting for resistance status and country (p > 0.05).

## RESULTS

In order to compare the varroa sizes in non-resistant and varroa-resistant honey bee colonies, we measured the dorsal shield area of mites collected from four European countries: Czech Republic, UK (Wales), Sweden and France. In each country, several colonies were sampled from at least one varroa-resistant apiary where bees had not received miticide treatment for several years, as well as from at least one control non-resistant apiary with regular miticide treatment. Resistance was achieved either through natural selection (UK) or through selective breeding for varroa-resistant traits (Czechia, France, Sweden). In total we analyzed over 4000 mites from nearly 180 colonies (Fig. 1).

**Fig. 1:**
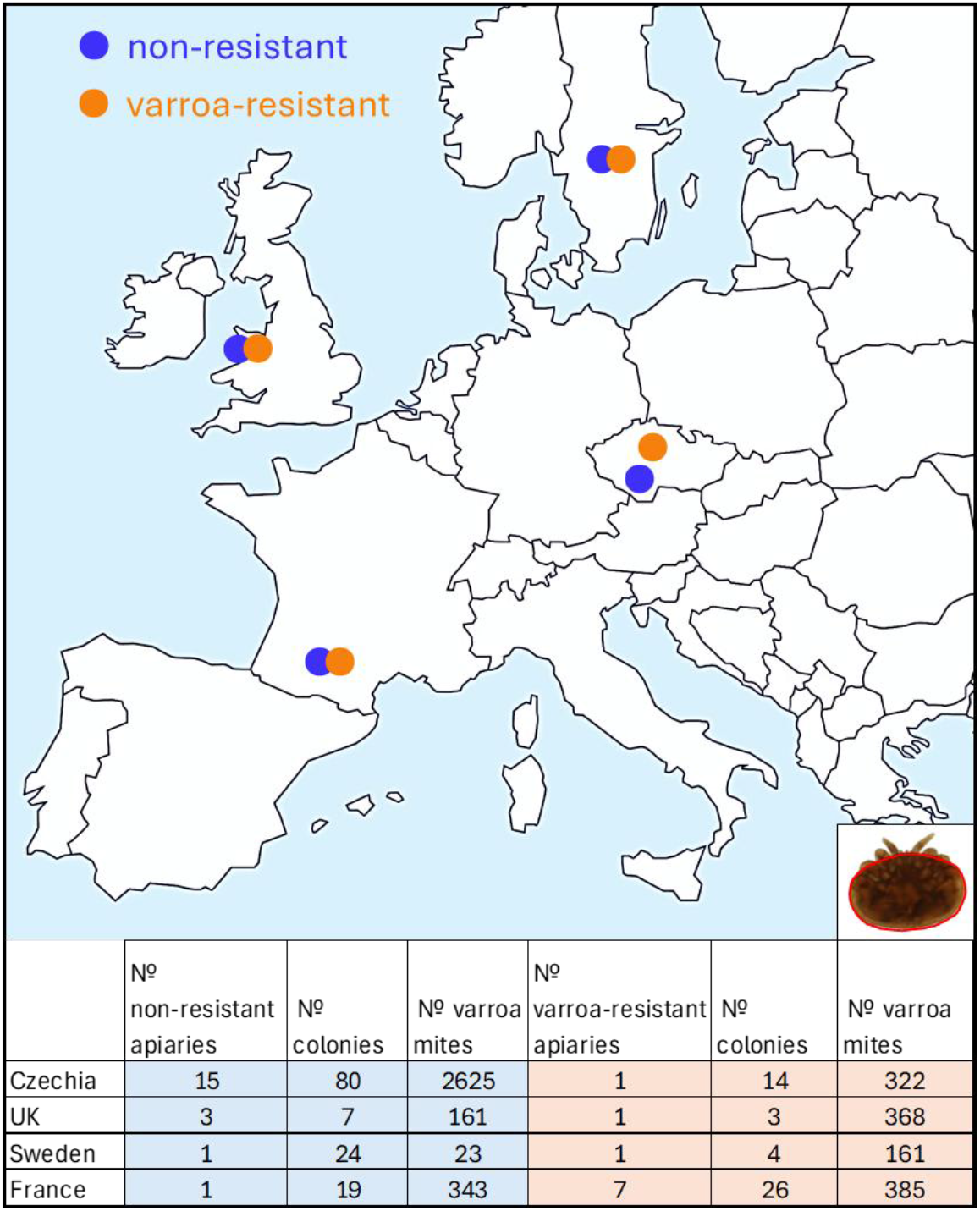
Geographical distribution of apiaries and the design of the study. The inset in the bottom right corner shows the analyzed area of the dorsal shield (marked in red). For GPS locations of all apiaries, beekeepers names, date of sample collections and type of hives used refer to table ST1.

In Czechia, robust data for mites from control non-resistant apiaries were collected, as varroa from 80 colonies in 15 apiaries was analyzed within two years. The median area of the mite dorsal shield across all apiaries was similar between the two years, 1.47 mm^2^ and 1.46 mm^2^ respectively (Fig. 2A and 2B). The median dorsal shield area of mites from varroa-resistant colonies was 1.39 mm^2^ (Fig. 2C), representing a statistically significant difference from mites in non-resistant colonies (Fig. 2D).

**Fig. 2:**
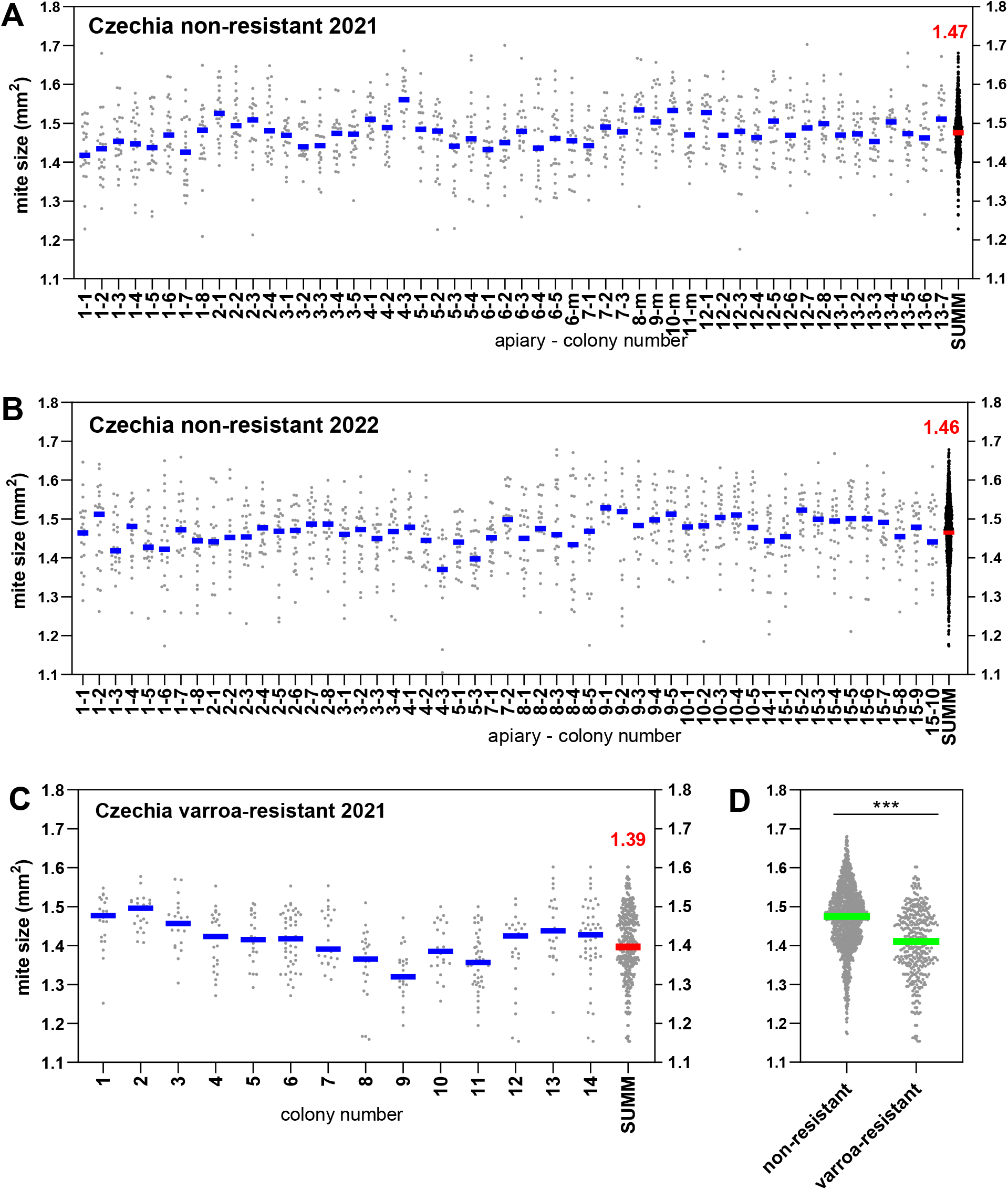
Varroa mite size in Czechia. (**A**) Mites from non-resistant, varroa-treated colonies collected in October 2021. The letter “m” in the legend to x-axis indicates a mixed sample containing mites from several colonies within the apiary. (**B**) Mites from non-resistant, varroa-treated colonies collected in October 2022. The apiary identifiers are the same in both years but the specific colony numbers do not match. (**C**) Mites from varroa-resistant, non-treated colonies collected in October 2021. (**D**) Summary of all Czech non-resistant and varroa-resistant mites collected in 2021 and 2022 (pooled data). Significance according to Student’s t-test (p<0.001). Each point represents dorsal shield size of an individual mite; horizontal line indicates the median value. The column with the red median line represents the combined data from all values shown within the graph. For list of apiaries refer to supplementary table ST1.

In the UK, the median area of the mite dorsal shield from control non-resistant colonies was 1.47 mm^2^ (Fig. 3A), similar to the control values in Czechia. Three colonies across one varroa-resistant apiary were sampled repeatedly during the season. Although the median values varied between the sampling points, the median values of all mites collected within the season in each colony were similar (1.36 mm^2^, 1.38 mm^2^ and 1.34 mm^2^, respectively, Fig. 3B). There was a statistically significant difference between the dorsal shield size of mites originating from the non-resistant and varroa-resistant colonies (Fig. 3C).

**Fig. 3:**
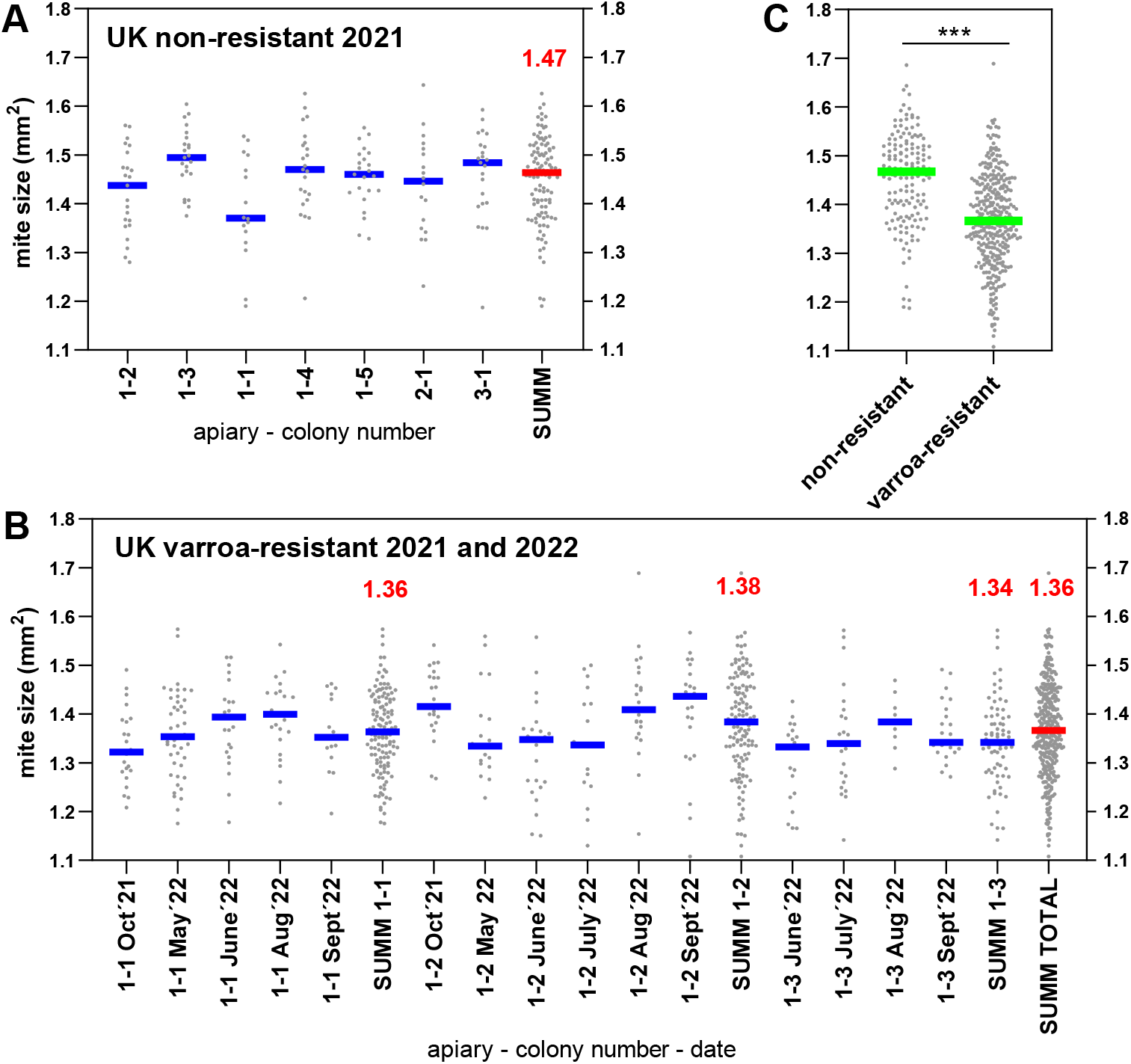
Varroa mite size in United Kingdom. (**A**) Size of varroa mites from non-resistant, varroa-treated colonies collected in September 2021. (**B**) Size of varroa mites from varroa-resistant, non-treated colonies of David Heaf collected on indicated dates in 2021 and 2022. (**C**) Summary of all UK non-resistant and varroa-resistant mites collected in 2021 and 2022 (pooled data). Significance according to Student’s t-test (p<0.001). Each point represents dorsal shield size of an individual mite; horizontal line indicates the median value. The column with the red median line represents the combined data from all values shown in the graph. For list of apiaries refer to supplementary table ST1.

In Sweden, only one control non-resistant apiary was available and the analyzed sample comprised mixed debris from twenty four colonies. The median mite dorsal shield area still remained similar to control values from other countries (1.47 mm^2^; Fig. 4A). The median dorsal shield area of mites from varroa-resistant colonies was 1.41 mm^2^ (Fig. 4B), representing a statistically significant difference from mites in non-resistant colonies (Fig. 4C).

**Fig. 4:**
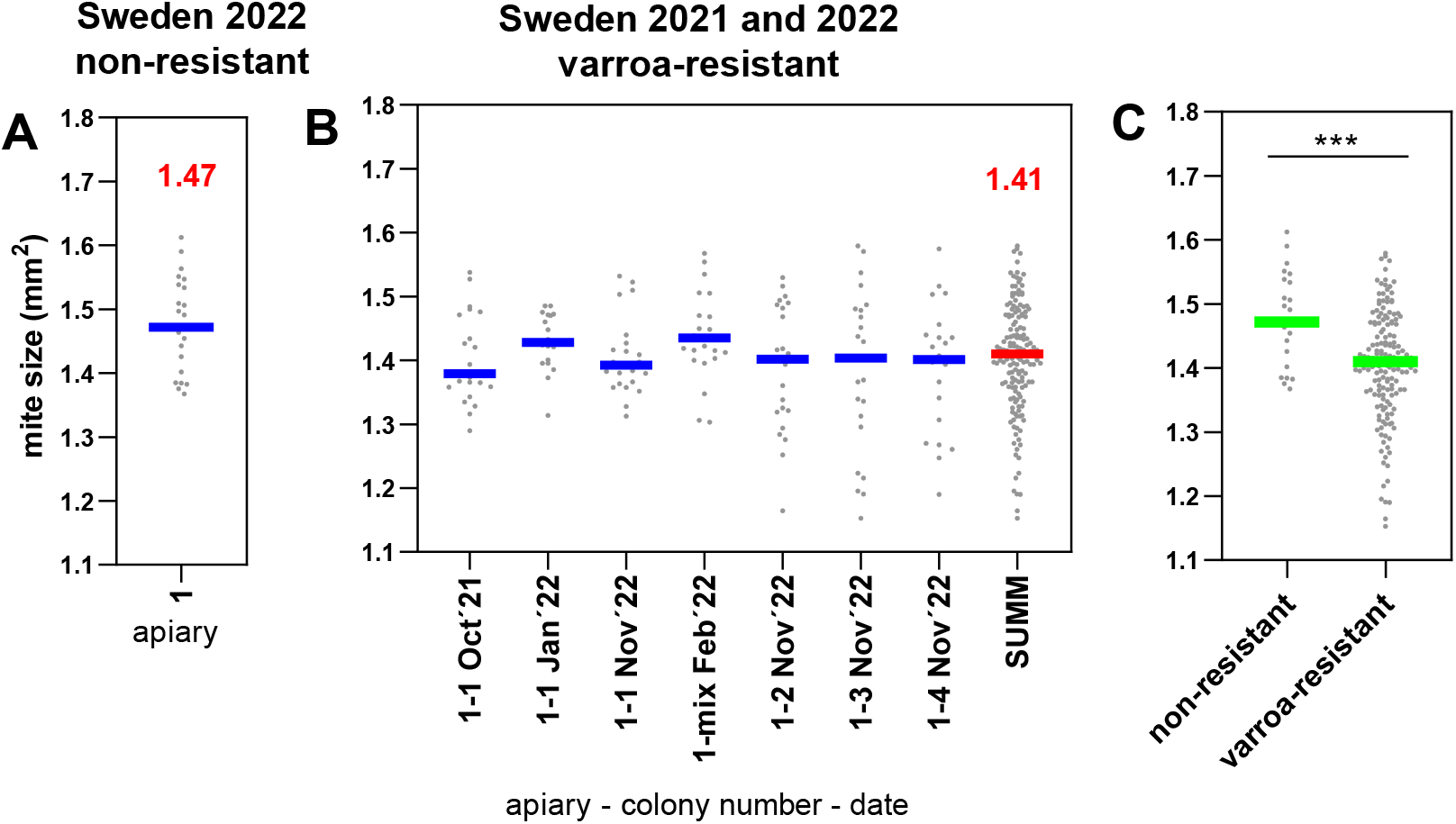
Varroa mite size in Sweden. (**A**) Size of varroa mites from non-resistant, varroa-treated apiary in October 2022. Mixed sample from 24 colonies. (**B**) Size of varroa mites from varroa-resistant, non-treated colonies of Eric Österlund collected on indicated dates in 2021 and 2022. (**C**) Summary of all Swedish non-resistant and varroa-resistant mites collected in 2021 and 2022 (pooled data). Significance according to Student’s t-test (p<0.001). Each point represents dorsal shield the size of an individual mite; horizontal line indicates the median value. The column with the red median line represents the combined data from all values shown in the graph. For list of apiaries refer to supplementary table ST1.

In France, we analyzed three groups of samples: (1) mites from varroa-resistant colonies not treated by miticides, (2) mites from varroa-resistant colonies that were treated with miticides and (3) mites from colonies where no intentional varroa-resistance selection was performed, the colonies were treated with miticides and they were located in a close proximity to the varroa-resistant apiaries from groups (1) and (2). The median dorsal shield size of mites from varroa-resistant, non-treated apiary was 1.35 mm^2^ (Fig. 5A). Regardless of miticide treatment, the mites from varroa-resistant colonies in group 2 were still small, with the median shield size of 1.34 mm^2^ (Fig. 5B). The mites from group 3 were originally thought to serve as controls, originating from presumably non-resistant, miticide treated colonies. However, we were not aware of its proximity to two varroa-resistant apiaries. The median size of mites from this apiary was 1.36 mm^2^ (Fig. 5C) suggesting that the close location of this apiary to the varroa-resistant apiaries lead to unintentional introduction of varroa-resistant traits (via queen mating with drones from the nearby varroa-resistant colonies) or to small mites introduction via robbing from varroa-resistant colonies. Mites from all three groups of samples were small and their dorsal shield sizes did not statistically differ (Fig. 5D).

**Fig. 5:**
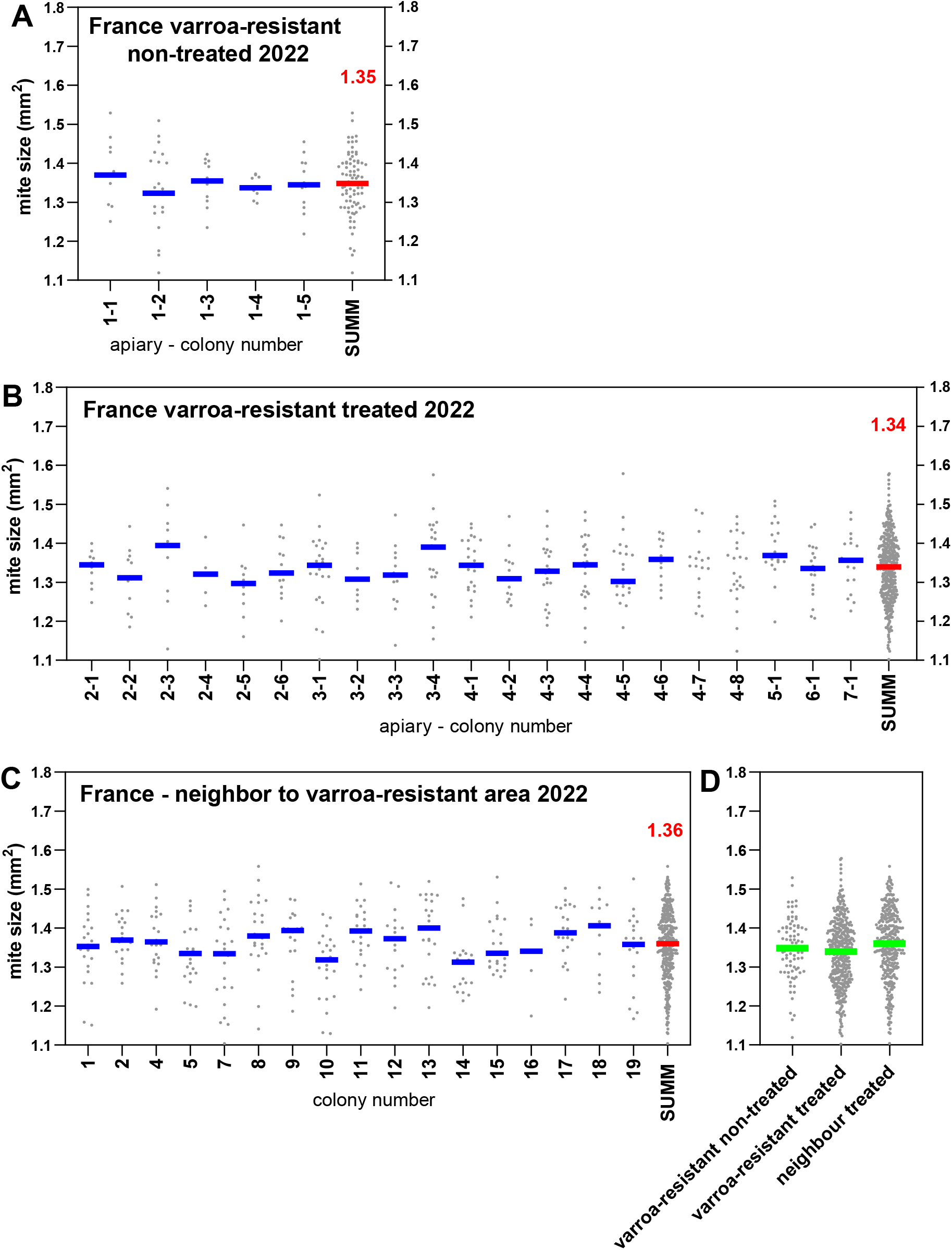
Varroa mite size in France. (**A**) Size of varroa mites from varroa-resistant, non-treated colonies of John Kefuss collected in September 2022. (**B**) Size of varroa mites from varroa-resistant but still varroa-treated colonies of Cyril Kefuss collected in September 2022. (**C**) Size of varroa mites collected from varroa-treated colonies of a neighboring beekeeper located 1.8 km and 3.72 km far from varroa-resistant apiaries. Samples collected in September 2022. (**D**) Summary of all French mites collected in 2022. Differences are not significant according to Student’s t-test. Each point represents dorsal shield size of an individual mite; horizontal line indicates the median value. The column with the red median line represents the combined data from all values shown in the graph. For list of apiaries refer to supplementary table ST1.

Taken together, dorsal shield sizes of mites from non-resistant honey bee colonies were highly consistent across countries, with median values of 1.47 mm^2^ (Fig. 6A). Although greater inter-country variability was observed in mites from varroa-resistant colonies, their dorsal shields were consistently smaller (median 1.37 mm^2^), representing an approximately 6.8% size reduction relative to mites from non-resistant colonies (Fig. 6B).

**Fig. 6:**
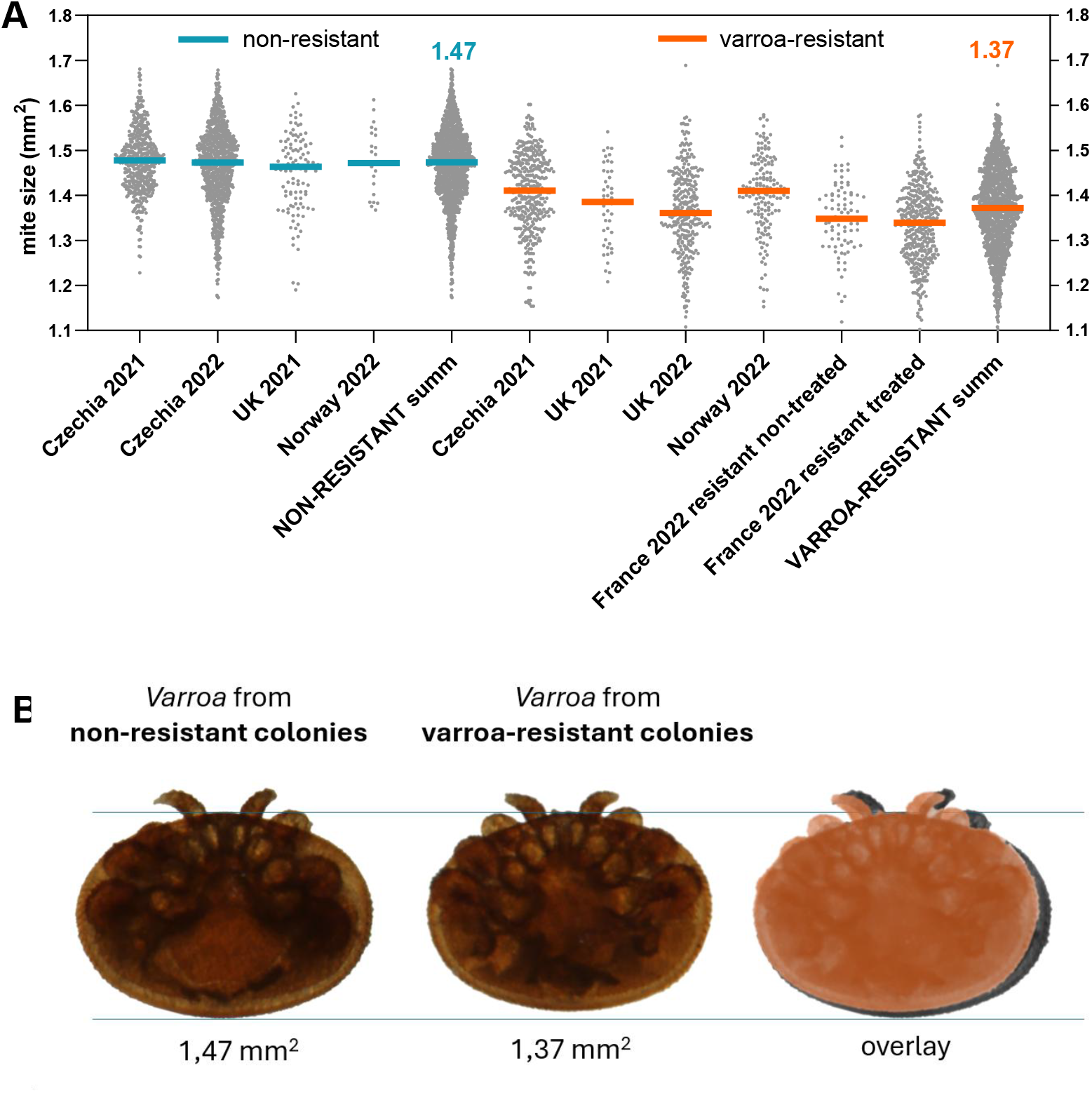
Summary of mite size data obtained from non-resistant and varroa-resistant colonies in Czechia, UK, Sweden and France. **(A)** Pooled data of all mites collected in indicated states and years. Each point represents dorsal shield size of an individual mite; horizontal line indicates the median value. Mites from resistant colonies are significantly smaller than those from susceptible colonies (linear mixed-effects model with colony as a random factor, p < 0.001) and the effect does not differ among countries (interaction test p = 0.36). **(B)** Representative images of average sized *Varroa destructor* mites from non-resistant and varroa-resistant honey bee colonies, with an overlay demonstrating a 6.8% difference in the size of the mites dorsal shields.

## DISCUSSION

### Reduced body size as a consistent pattern across Europe

Our data demonstrate that *Varroa destructor* mites originating from varroa-resistant honey bee populations are consistently smaller than mites from non-resistant colonies across geographically distinct European regions. Despite variation in location, bee genetic origin and management practices, the median dorsal shield area of mites from non-resistant colonies remained remarkably stable (1.47 mm^2^). Mites from varroa-resistant colonies exhibited a significant reduction in body size (median 1.37 mm^2^), corresponding to an approximately 6.8% difference.

The consistency of values across countries strengthens the biological relevance of the observed size reduction. Statistical analyses confirmed that the smaller body size of mites associated with resistant colonies depended on the resistance status of the colony rather than on country of origin or miticide treatment. These results indicate that the reduced size of mites in varroa-resistant colonies represents a host associated effect.

### Morphological shift as a potential consequence of host imposed pressure

Body size in invertebrates is an integrative trait reflecting nutritional conditions, developmental timing and reproductive allocation [26]. Because *Varroa destructor* completes its reproductive cycle within capped brood cells, its development is tightly coupled with the conditions mediated by the host. Varroa resistant honey bee colonies display characteristic host behavior that lead to reduced overall reproductive success of the varroa mite within the colony [10; 11] and to better colony survival. This includes varroa-sensitive hygiene, cell recapping or grooming. These host behavioral traits may alter the mites feeding frequency or efficiency on developing honey bee pupae that can lead to nutritional limitation, affecting the mite body size and its reproductive success.

Disbalance in hormones promoting growth could also result in reduced mite body size. Ecdysone is a hormone controlling metamorphosis and affecting organismal growth [27]. It is also crucial for the start of mite oogenesis and the transition between the young mite developmental stages [27; 28]. Varroa can not synthesize this hormone on its own but relies on its supply from the host pupa [29]. Interestingly, mutations in the ecdysone synthesis pathway in honey bees have been linked to varroa resistance [29]. Changes in the timing of ecdysone expression in honey bee pupae from varroa-resistant colonies could therefore contribute to the small varroa size and represent mite developmental constrain under host pressure.

Another host-derived factor that could influence mite size during development is the temperature of the honey bee brood nest. The well established temperature–size rule states that ectotherms reared at higher temperatures typically mature at smaller body sizes [30]. This rule applies also to the *Acari* family, as the spider mite *Tetranychus ludeni* produces smaller adults at higher temperatures [31]. Honey bees maintain their brood nest temperature close to 35**°**C [32] but variations within the range of 33-36 °C occur[33]. Maintaining brood nest temperatures at the upper end of this range in varroa-resistant colonies could potentially result in smaller body size of varroa mites developing within the brood. However, a comparative study examining long term brood nest temperature profiles in varroa-resistant versus susceptible colonies is currently lacking.

### Potential effect of small body size on varroa reproduction

Body size in parasitic arthropods is often associated with fecundity, dispersal capacity and feeding efficiency. In some species of predatory mites (e.g., *Phytoseiulus persimilis*), early food limitation during female development leads to smaller adults with reduced fecundity, lower maternal survival probability, decreased attractiveness for males and a reduced number and size of eggs compared to standard size females [34]. Similarly in spider mites (*Tetranychus* spp.), multiple studies have reported a positive correlation between body size and fecundity, where larger females lay more eggs [35]. Although the effect of varroa body size on fecundity or survival has not been studied, the above mentioned examples clearly indicate fitness costs associated with smaller body size in the *Acari* family.

If smaller varroa mites produce fewer offspring and exhibit altered reproductive success, reduced body size may contribute to the observed slow mite reproduction phenotype frequently observed in varroa-resistant colonies [10] and lead to stabilization of host-parasite equilibrium observed in naturally surviving populations.

### Effect of miticide treatment, type of comb and type of hive

As all the non-resistant colonies in the study were treated with miticides, one could speculate that miticide treatment might be responsible for the larger mite size. The data from France provide an important insight about this question as mites from French varroa-resistant colonies that were treated with amitraz remained small, comparable in size to mites from untreated resistant colonies. Accordingly, there was no statistically significant effect of varroa treatment on mite size in our data wehn tested using a linear mixed-effects model. Smaller mites were observed in colonies living on various types of combs – freely built combs (built by the bees without the use of a wax foundation provided by the beekeeper), combs built from wax foundations of regular size or from wax foundations of smaller size. The types of hive also differed substantially amongst the countries and individual apiaries, showing no obvious pattern related to the mite sizes.

These findings suggest that miticide treatment, the type of comb or the type of hive used for beekeeping do not determine mite body size. Instead, the host varroa-resistant phenotype arising from long term selection appears to be the dominant factor.

### Sampling strategy to select varroa-resistant colonies based on varroa size

Quantifying the dorsal shield area of *Varroa destructor* provides several methodological advantages over measuring the mite body weight or other morphological parameters. It is a stable structural trait that is not influenced by gut contents or by the presence or absence of an egg in the female. It is also not affected by sample storage conditions, allowing retrospective analysis of samples. The dorsal shield area is easy to distinguish and digital image analysis leads to high reproducibility. Therefore, dorsal shield size may represent a robust morphological parameter suitable for comparative studies across populations. From this perspective, measuring the dorsal shield area could serve as a new practical marker for selecting varroa-resistant colonies, together with established criteria such as mite fertility in brood [36] or the PIN test [37].

In Sweden, seasonal differences in mite dorsal shield size were observed at individual time points that likely reflect morphological variations of varroa during the year [38]. However, when data were pooled across the entire season, mites from resistant colonies showed similar median sizes. This single time point sampling may therefore not accurately reflect the long term phenotype of the parasite population. Seasonal pooling appears more appropriate when assessing stable morphological trends and this should be remembered as an important methodological consideration when using mite size as a selection marker for varroa-resistant honey bee colonies.

### Future research directions

A limitation of the present study is that the varroa size was not correlated with the mite fecundity or other physiological parameters. Future studies should correlate the varroa size with its reproductive success, longevity and potentially also with its viral loads. Such integrative approaches would clarify whether reduced body size correlates with reduced parasite fitness.

## Conclusions

Across four European countries, *Varroa destructor* mites associated with varroa-resistant honey bee colonies consistently exhibited reduced dorsal shield size compared with mites from non-resistant colonies. This pattern was reproducible across distinct geographical regions and it was independent of the honey bees genetic background, miticide treatment or colony management practices. These findings suggest that long term host resistance influences parasite morphology and highlight dorsal shield size as a potentially informative new phenotypic marker of varroa-resistance.

## Acknowledgement

We thank to Jiří Marx, John Kefuss, Cyril Kefuss, Victor Kohut, Eric Österlund, A.Andersson, David Heaf, Jonathan Garratt and all the other beekeepers who kindly provided debris samples from their honey bee colonies. This study was supported by the project reg. no. CZ.02.01.01/00/23_021/0009529 (iBEEIntelligent Beekeeping: Modern Biotechnologies for Healthy Bees), co-funded by the European Union and by the Czech Science Foundation (25-16047S).

## FIGURE LEGEND

Supplementary table ST1: List of apiaries, their GPS locations, beekeepers names, types of hives and dates of mite collection.

